# Red Knot Stopover Population Size and Migration Ecology at Delaware Bay, USA, 2021

**DOI:** 10.1101/2022.03.23.485371

**Authors:** J. E. Lyons

## Abstract

Red Knots (*Calidris canutus rufa*) stop at Delaware Bay during northward migration to feed on eggs of horseshoe crabs (*Limulus polyphemus*). In the late 1990s and early 2000s, the number of Red Knots found at Delaware Bay dramatically declined from ~50,000 to ~13,000. Horseshoe crabs have been harvested for use as bait in eel and whelk fisheries since at least 1990, and some avian conservation biologists hypothesized that harvest levels in the 1990s prevented sufficient refueling for successful migration to the breeding grounds, nesting, and survival for the remainder of the annual cycle. Since 2013, the harvest of horseshoe crabs in the Delaware Bay region has been managed using an Adaptive Resource Management (ARM) framework. The objective of the ARM framework is to manage sustainable harvest of Delaware Bay horseshoe crabs while maintaining ecosystem integrity and supporting Red Knot recovery with adequate stopover habitat for Red Knots and other migrating shorebirds. For annual harvest recommendations, the ARM framework requires annual estimates of horseshoe crab population size and the Red Knot stopover population. We conducted a mark-recapture-resight investigation to estimate the passage population of Red Knots at Delaware Bay in 2021. We used a Bayesian analysis of a Jolly-Seber model, which accounts for turnover in the population and the probability of detection during surveys. The 2021 Red Knot mark-resight dataset included a total of 1,591 individual birds that were recorded at least one during mark-resight surveys at Delaware Bay in 2021. The passage population size in 2021 was estimated at 42,271 (95% credible interval: 35,948 – 55,210). Like 2020, the 2021 population estimate is slightly lower than the 2018 and 2019 estimates. The 2021 population size estimate will inform decision making for harvest recommendations in the next management cycle.

## 2 Background

Red Knots (*Calidris canutus rufa*) stop at Delaware Bay during northward migration to feed on eggs of horseshoe crabs (*Limulus polyphemus*). The northward migration of *C. c. rufa* coincides with the spawning of horseshoe crabs, whose eggs are the perfect food for a migrating Red Knot (Karpanty et al. 2006, Haramis et al. 2007). Horseshoe crabs are therefore an important food resource for Red Knots as well as other shorebirds at Delaware Bay.

Horseshoe crabs have been harvested since at least 1990 for use as bait in American eel (*Anguilla rostrata*) and whelk (*Busycon*) fisheries (Kreamer and Michels 2009). In the late 1990s and early 2000s the number of Red Knots found at Delaware Bay declined dramatically from ~50,000 to ~13,000 (Niles et al. 2008). At the same time the number of horseshoe crabs harvested also declined and avian conservation biologists hypothesized that unregulated harvest of horseshoe crabs from Delaware Bay in the 1990s prevented sufficient refueling during stopover for successful migration to the breeding grounds, nesting, and survival for the remainder of the annual cycle (McGowan et al. 2011).

The harvest of horseshoe crabs in the Delaware Bay region has been managed by the Atlantic States Marine Fisheries Commission (ASMFC) since 2012 using an Adaptive Resource Management (ARM) framework (McGowan et al. 2015b). The ARM framework was designed to constrain the harvest so that number of spawning crabs would not limit the number of Red Knots stopping at Delaware Bay during migration. This management framework to achieve multiple objectives requires an estimate each year of both the crab population and the Red Knot stopover population size to inform harvest recommendations (McGowan et al. 2015a). We have estimated the stopover population size using mark-resight data on individually-marked birds and a Jolly-Seber model for open populations since 2011.

## 3 Methods

Red knots have been individually marked at Delaware Bay and other locations in the Western Hemisphere with engraved leg flags since 2003; each leg flag is engraved with a unique 3-character alphanumeric code (Clark et al. 2005). Mark-resight data (i.e., sight records of individually-marked birds and counts of marked and unmarked birds) were collected on the Delaware and New Jersey shores of Delaware Bay in 2021 according to the methods for mark-resight investigations of Red Knots at Delaware Bay (Lyons 2016). This protocol has been used at Delaware Bay since 2011.

Surveys to locate leg-flagged birds were conducted on each beach in 2021, every three days in May and June according to the sampling plan (Table 1). During these resighting surveys, agency staff and volunteers surveyed the entire beach and recorded as many alphanumeric combinations as possible.

**Table 1.** Dates for mark-resight survey periods (3-day sampling occasions) at Delaware Bay. Survey period 10 was not used in 2021 because the mark-resight data were sparse in this period.

| Survey period | Dates | Survey period | Dates |
| --- | --- | --- | --- |
| 1 | ≤10 May | 6 | 23-25 May |
| 2 | 11-13 May | 7 | 26-28 May |
| 3 | 14-16 May | 8 | 29-31 May |
| 4 | 17-19 May | 9 | 1-3 June |
| 5 | 20-22 May | 10 | 4-6 June |

As in previous years, all flag resightings were validated with physical capture and banding data available in the data repository at http://www.bandedbirds.org/. Resightings without a corresponding record of physical capture and banding (i.e., “misread” errors) were not included in the analysis. However, banding data from Argentina are not available for validation purposes in bandedbirds.org; therefore, all resightings of orange engraved flags were included in the analysis without validation using banding data. We also omitted resightings of 12 flagged individuals in 2021 whose flag codes were previously accidentally deployed in both New Jersey and South Carolina (Amanda Dey, New Jersey Division of Fish and Wildlife, pers. comm., 31 May 2017) because it is not possible to confirm individual identity in this case. Section 4 “Summary of Mark-resight and Count Data Collected in 2021” describes additional quality control procedures and the potential for other types of errors in the mark-resight dataset.

While searching for birds marked with engraved leg flags, observers also periodically used a scan sampling technique to count marked and unmarked birds in randomly selected portions of Red Knot flocks (Lyons 2016).

To estimate stopover population size, we used the methods of Lyons et al. (2016) to analyze 1) the mark-resight data (flag codes), and 2) data from the scan samples of the marked:unmarked ratio. Lyons et al. (2016) rely on the “superpopulation” approach developed by Crosbie and Manly (1985) and Schwarz and Arnason (1996). The superpopulation is defined as the total number of birds present in the study area on at least one of the sampling occasions over the entire study, i.e., the total number of birds present in the study area at any time between the first and last sampling occasions (Nichols and Kaiser 1999). In this superpopulation approach, passage population size is estimated each year using the Jolly-Seber model for open populations, which accounts for the flow-through nature of migration areas and probability of detection during surveys.

In our analyses for Delaware Bay, the days of the migration season were aggregated into 3-day sampling periods (a total of 10 sample periods possible each season, Table 1). Data were aggregated to 3-day periods because this is the amount of time necessary to complete mark-resight surveys on all beaches in the study (a summary of the mark-resight data from 2021 is provided in Appendix 1).

With the mark-resight superpopulation approach, we first estimated the number of birds that were carrying leg flags, and then adjusted this number to account for unmarked birds using the estimated proportion of the population with flags. The estimated proportion with leg flags is thus an important statistic. We used the scan sample data (i.e., the counts of marked birds and the number checked for marks) and a binomial model to estimate the proportion of the population that is marked. To account for the random nature of arrival of marked birds in the bay and the addition of new marks during the season, we implemented the binomial model as a generalized linear mixed model with a random effect for the sampling period. More detailed methods are provided in Lyons et al. (2016) and Appendix 2.

## 4 Summary of Mark-resight and Count Data Collected in 2021

### Mark-resight encounter data

The 2021 Red Knot mark-resight dataset included a total of 1,591 individual birds that were recorded at least one during mark-resight surveys at Delaware Bay in 2021; these birds were originally captured and banded with leg flags in five different countries (Table 2). This total is remarkably close to the 2020 total detected at Delaware Bay: 1,587 individual birds were recorded in 2020 (Table 2). **Approximately the same number of flagged Red Knots were detected at Delaware Bay in 2020 and 2021.**

**Table 2.** Number of Red Knot (*C. c. rufa*) flags detected in 2021 by banding location (flag color).

|  | ***No. flagged individuals detected*** |  |  |
| --- | --- | --- | --- |
| ***Banding location (flag color)*** | ***2019*** | ***2020*** | ***2021*** |
| U.S. (lime green) | 2,368 | 1,255 | 1,292 |
| U.S. (dark green) | 351 | 161 | 118 |
| Argentina (orange) | 216 | 89 | 81 |
| Canada (white) | 156 | 52 | 78 |
| Brazil (dark blue) | 35 | 21 | 17 |
| Chile (red) | 10 | 9 | 5 |
| Total | 3,136 | 1,587 | 1,591 |

There was sufficient data for analysis in all 10 sampling periods in 2021 (≤10 May to 6 June; Table 1). In some years, including 2020, the analysis was restricted to periods 1-9 (≤10 May to 3 June) because data beyond 3 June were sparse.

While the number of birds detected in 2021 was similar to the number detected in 2020, this number of individuals resighted within a season is lower than recent (pre-COVID-19) years given the limited use of volunteers for safety reasons. The number of marked birds detected and available for analysis in 2021 was approximately 48% lower than the number in the 2019 analysis (n = 3,072 birds) and 58% lower than the number detected and used for analysis in 2018 (n = 3,820).

One assumption of the mark-resight approach is that individual identity of marked birds is recorded without error (see Lyons 2016 for discussion of all model assumptions). As noted above, some field-recording errors are evident when sight records are compared to physical capture records available from bandedbirds.org. Again, any engraved flag reported by observers that does not have a corresponding record of physical capture is omitted. Field observers submitted 3,792 resightings in 2021; 50 were not valid (i.e., no corresponding banding data), for an overall misread read of 1.3%. (In 2020, 3,364 resightings were submitted and 100 [2.9%] were not valid.) These invalid resightings were removed before analysis, but a second type of “false positive” is still possible, i.e., false positive detection of flags that were deployed prior to 2021 but were not in fact present at Delaware Bay in 2021. It is not possible to identify this second type of false positive with banding data validation or other quality assurance/quality control methods.

### Marked-ratio data

In 2021, 564 marked ratio scan samples were collected: 297 and 267 samples in Delaware and New Jersey, respectively (Appendix 3). Last year in 2020, 734 marked-ratio scan samples were collected: 376 samples in Delaware and 358 in New Jersey.

### Aerial and ground count data

Aerial surveys were conducted on 23 and 27 May 2021 (Table 3; data provided by A. Dey, New Jersey Division of Fish and Wildlife, Endangered and Nongame Species Program). Ground and boat surveys were conducted twice in New Jersey (on 23 and 27 May) but only once in Delaware (on 23 May; Table 3).

**Table 3.** Number of Red Knots detected during aerial and ground surveys of Delaware Bay in 2021. Data provided by A. Dey, New Jersey Division of Fish and Wildlife, Endangered and Nongame Species Program.

|  | Delaware | New Jersey | Total |
| --- | --- | --- | --- |
| **Aerial/Ground Surveys** |  |  |  |
| 23 May 2021 | 1,123* | 5,012 | 6,131 |
| 27 May 2021 | 895 | 5,985 | 6,880 |
| **Ground/Boat Surveys** |  |  |  |
| 23 May 2021 | 1,123 | 3,651 | 4,774 |
| 27 May 2021 | — | 5,618 | 5,618 |

\* Delaware ground survey total from 23 May (1,123 birds) used here rather than the aerial count of Delaware on the same day because the aerial count was lower than the corresponding ground count.
“—” = no data; ground survey was not conducted in Delaware on 27 May.

## 5 Summary of 2021 Migration

The pattern of arrivals at Delaware Bay in 2021 suggests a slow start to the migration season, with few birds arriving before 18 May. A large wave of arrivals occurred on or about 21 May: approximately 35% of the total 2021 stopover population arrived close to 21 May (Fig. 1a). The number of birds arriving in the following period, about 24 May, was low, but there was a small number of late arrivals around 27-31 May (approximately 21% of the stopover population). Thus in 2021, it appears there was one large wave of arrivals near the middle of the season and relatively small fractions arriving in the other the sampling periods before and after the peak of arrivals around 21 May.

**Figure 1.**
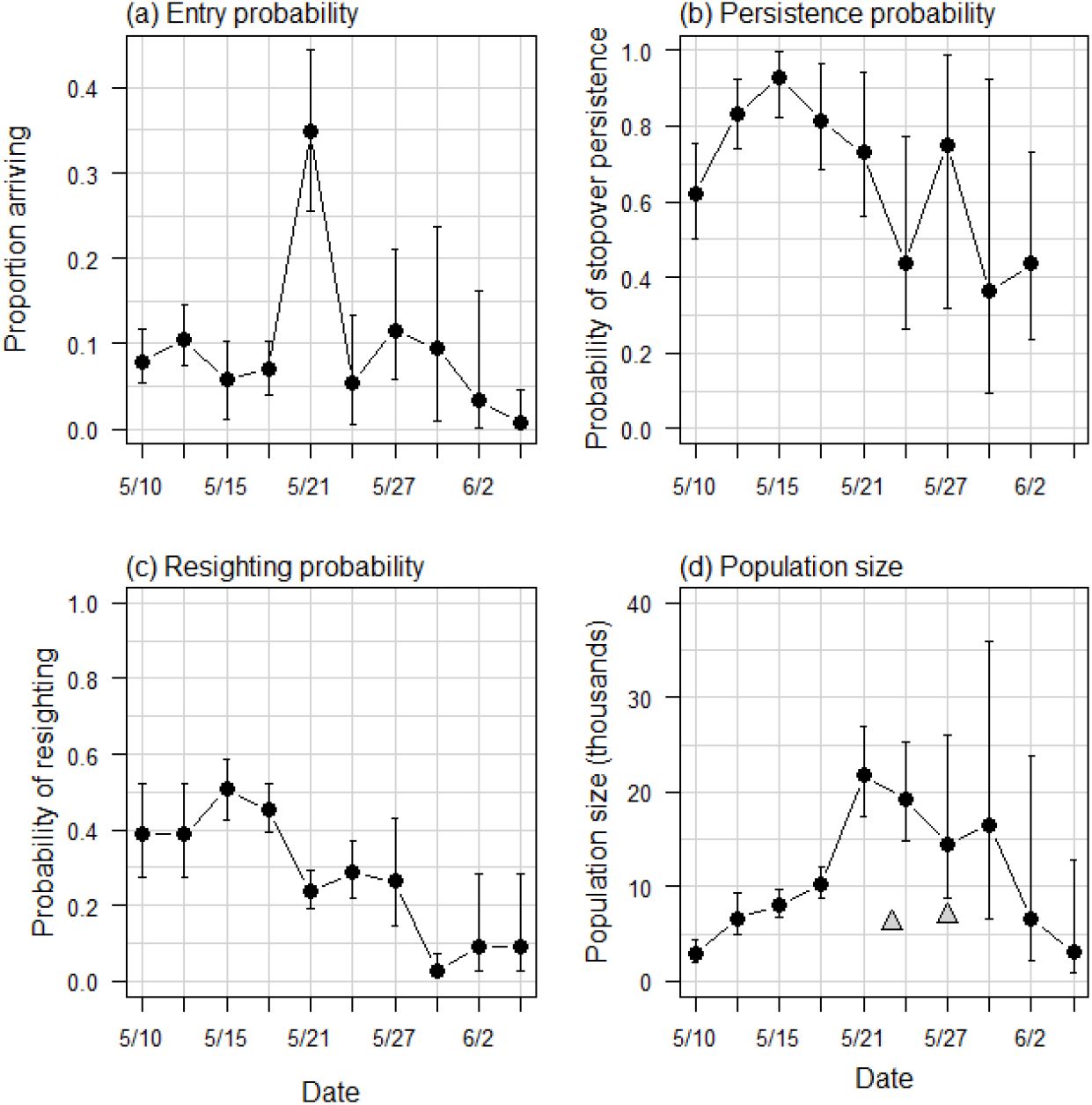
Estimated Jolly-Seber (JS) model parameters from a mark-resight study of Red Knots at Delaware Bay in 2021: (a) proportion of stopover population arriving at Delaware Bay, (b) stopover persistence, (c) probability of resighting, and (d) time-specific stopover population size. Dates on the x-axis represent sampling occasions (3-day survey periods). Triangles in (d) are total counts conducted on 23 (aerial count of NJ; ground count of DE) and 27 May (aerial count for both NJ and DE) 2021.

**Figure 2.**
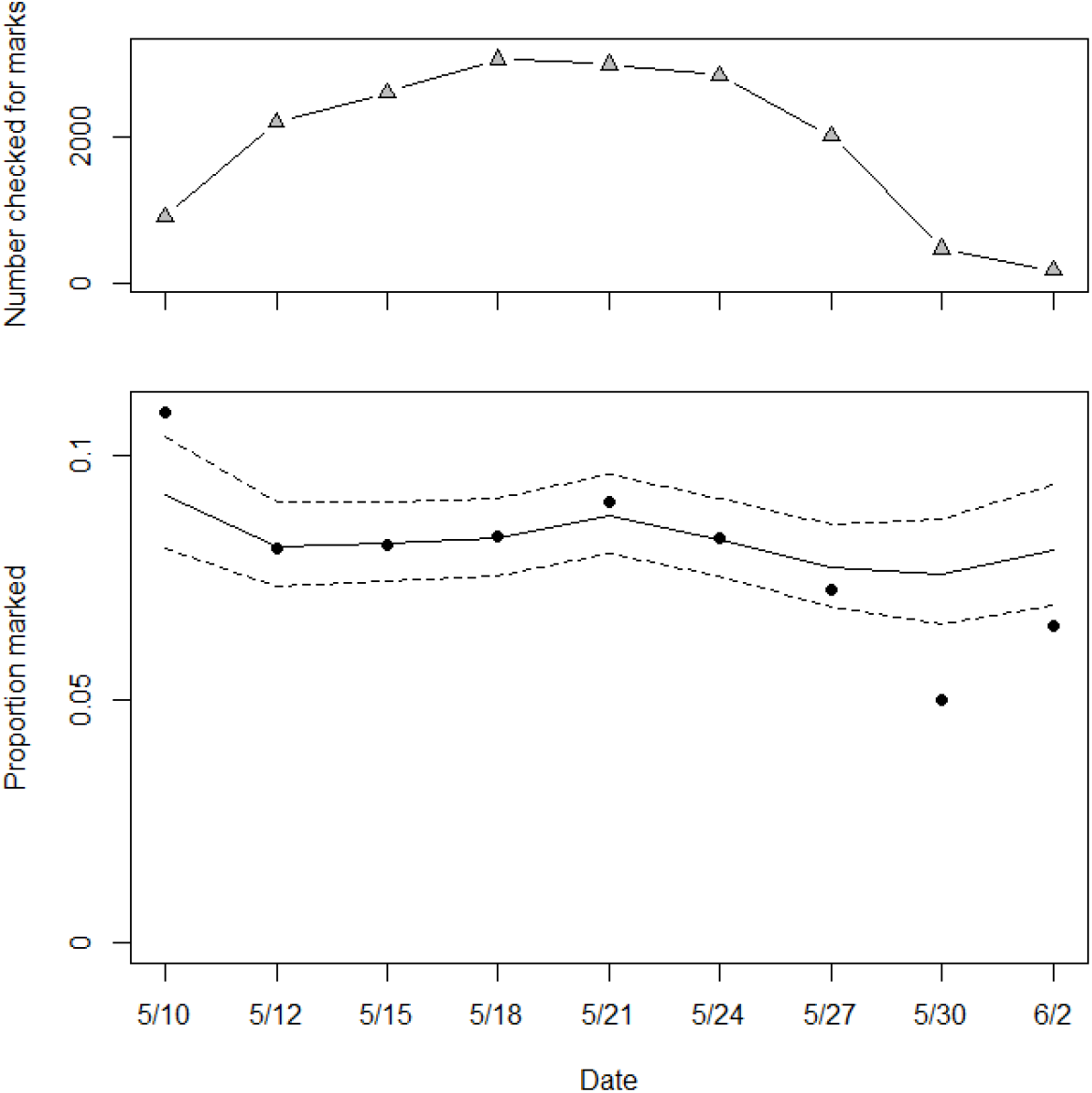
Estimated proportion of the Delaware Bay stopover population carrying leg flags in 2021. The marked proportion was estimated from marked-ratio scan samples for each 3-day sampling period. The dates for the sampling periods are shown in Table 1. The upper panel shows the sample size (number scanned, i.e., checked for marks) for each sample period. The bottom panel shows the estimated proportion marked at each sample occasion, which was estimated with the generalized linear mixed model described in Appendix 2. Solid and dashed lines are estimated median proportion marked and 95% credible interval; filled circles show (number with marks/number scanned).

Stopover persistence is the probability that a bird present at Delaware Bay during sampling period *i* is present at sampling period *i* + 1. In 2021, stopover persistence started off relatively low (0.6), which is unusual for this time of year (Fig 1b). Often the early-arriving birds remain in the study area with little turnover in the population (but see 2020), but in 2021 there was substantial turnover early in the season. Stopover persistence peaked around 15 May and declined steadily after that until 27 May (Fig 1b). The steady decrease in stopover persistence during 15-24 May suggested a high degree of turnover and shorter stopovers than most years. There was a spike in stopover persistence around 27 May (Fig. 1b), during which turnover slowed briefly, but otherwise, stopover persistence declined steadily from 15 May until the end of the season. That is, turnover was high and increasing from 15 May on, suggesting shorter stays in 2021 than in most other years.

Following Lyons et al. (2016), we used the Jolly-Seber model to estimate stopover duration. In 2021, estimated average stopover duration was 10.3 days (95% credible interval 9.0 – 12.1 days). This stopover duration estimate is slightly shorter than 2020 (10.7 days [9.9 – 11.7]) and shorter than 2019 (12.1 days). This method of estimating stopover duration provides a coarse measure in our Delaware Bay study, however, because it is based on the number of sampling periods that a bird remained in the study area. For our Delaware Bay analysis, sampling periods are 3 days in which the data are aggregated (Table 1). To estimate stopover duration at Delaware Bay with this method, we first estimate the number of sampling periods that each bird remained in the study area and then multiply this by 3 (the number of days in each period) to estimate stopover duration in days. The resolution of the estimate is thus limited by the resolution of the time step in the mark-recapture model.

Probability of resighting in 2021 was relatively high early in the season, approximately 40-50% until around 18 May (Fig 1c). Between 21-27 May, probability of resighting was lower, around 25%. At the end of the season, after 27 May, probability of resighting was lower still, especially the 3-day period around 31 May. Around 31 May, the probability of resighting was close to zero, which is unusual for the mark-resight work at Delaware Bay (Fig 1c). Resighting probability increased slightly during 1-6 June to levels more typical for this time of year.

In 2021, 8.2% of the stopover population carried engraved leg flags (95% CI, 7.0% −9.1%). This is slightly lower than the 2020 estimate (9.6% with leg flags [95% CI 8.8 - 10.3%]).

## 6 Stopover Population Estimation

The passage population size in 2021 was estimated at 42,271 (95% credible interval: 35,948 – 55,210). This superpopulation estimate accounts for turnover in the population and probability of detection. The 2021 stopover population estimate is similar to the 2020 stopover population size estimate (given wide confidence intervals in both years), 40,444 (33,627 - 49,966), and slightly lower than the 2018-2019 estimates (Table 4).

**Table 4.** Red Knot stopover (passage) population estimate using mark-resight methods compared to peak-count index using aerial- or ground-survey methods at Delaware Bay. The mark-resight estimate of stopover (passage) population accounts for population turnover during migration; peak-count index, a single count on a single day, does not account for turnover.

| Year | Stopover population<sup>a</sup><br>(mark-resight $N^*$) | 95% CI<br>Stopover pop-<br>ulation $N^*$ | Peak-count index<br>[aerial (A) or<br>ground (G)] |
| --- | --- | --- | --- |
| 2011 | 43,570 | (40,880 – 46,570) | 12,804 (A)<sup>b</sup> |
| 2012 | 44,100 | (41,860 – 46,790) | 25,458 (G)<sup>c</sup> |
| 2013 | 48,955 | (39,119 – 63,130) | 25,596 (A)<sup>d</sup> |
| 2014 | 44,010 | (41,900 – 46,310) | 24,980 (A)<sup>c</sup> |
| 2015 | 60,727 | (55,568 – 68,732) | 24,890 (A)<sup>c</sup> |
| 2016 | 47,254 | (44,873 – 50,574) | 21,128 (A)<sup>b</sup> |
| 2017 | 49,405<sup>e</sup> | (46,368 – 53,109) | 17,969 (A)<sup>f</sup> |
| 2018 | 45,221 | (42,568 – 49,508) | 32,930 (A)<sup>b</sup> |
| 2019 | 45,133 | (42,269 – 48,393) | 30,880 (A)<sup>g</sup> |
| 2020 | 40,444 | (33,627 – 49,966) | 19,397 (G)<sup>c</sup> |
| 2021 | 42,271 | (35,948 – 55,210) | 6,880 (A)<sup>h</sup> |

<sup>a</sup> passage population estimate for entire season, including population turnover
<sup>b</sup> 23 May
<sup>c</sup> 24 May
<sup>d</sup> 28 May
<sup>e</sup> Data management procedures to reduce bias from recording errors in the field; data from observers with greater than average misread rate were not included in the analysis.
<sup>f</sup> 26 May
<sup>g</sup> 22 May
<sup>h</sup> 27 May

Like 2020, the 2021 population estimate is slightly lower than the 2018 and 2019 estimates (Table 4) and the confidence interval is wider. The uncertainty in the population estimate and wide confidence intervals are due in part to the low probability of resighting for many of the sampling periods during 2020-2021 compared to other years (early 2021 notwithstanding). The time-specific stopover population estimates in 2021 increased steadily from the beginning of the season and peaked around 21 May (21,846 birds; Fig. 1d), corresponding to the large influx of arrivals at this time (Fig. 1a). Time-specific estimates declined steadily from 21 May until 6 June (Fig. 1d). The relatively high degree of uncertainty (wide confidence interval) in the estimate for the 30 May period reflects the low probability of resighting at this time (Fig. 1c).

## 1 Acknowledgments

We thank the many volunteers in Delaware and New Jersey who collected mark-resight data in 2021. We are grateful to Henrietta Bellman (Delaware Division of Fish and Wildlife), Amanda Dey (New Jersey Division of Fish and Wildlife), Lena Usyk (bandedbirds.org), and numerous volunteers in Delaware and New Jersey for data entry and data management. We are grateful to David Kazyak and William Link for helpful peer-reviews as part of U.S. Geological Survey Fundamental Science Practices. Any use of trade, firm, or product names is for descriptive purposes only and does not imply endorsement by the U.S. Government.

## Appendix 1. Summary of 2021 mark-resight data (“m-array”). NR = never resighted

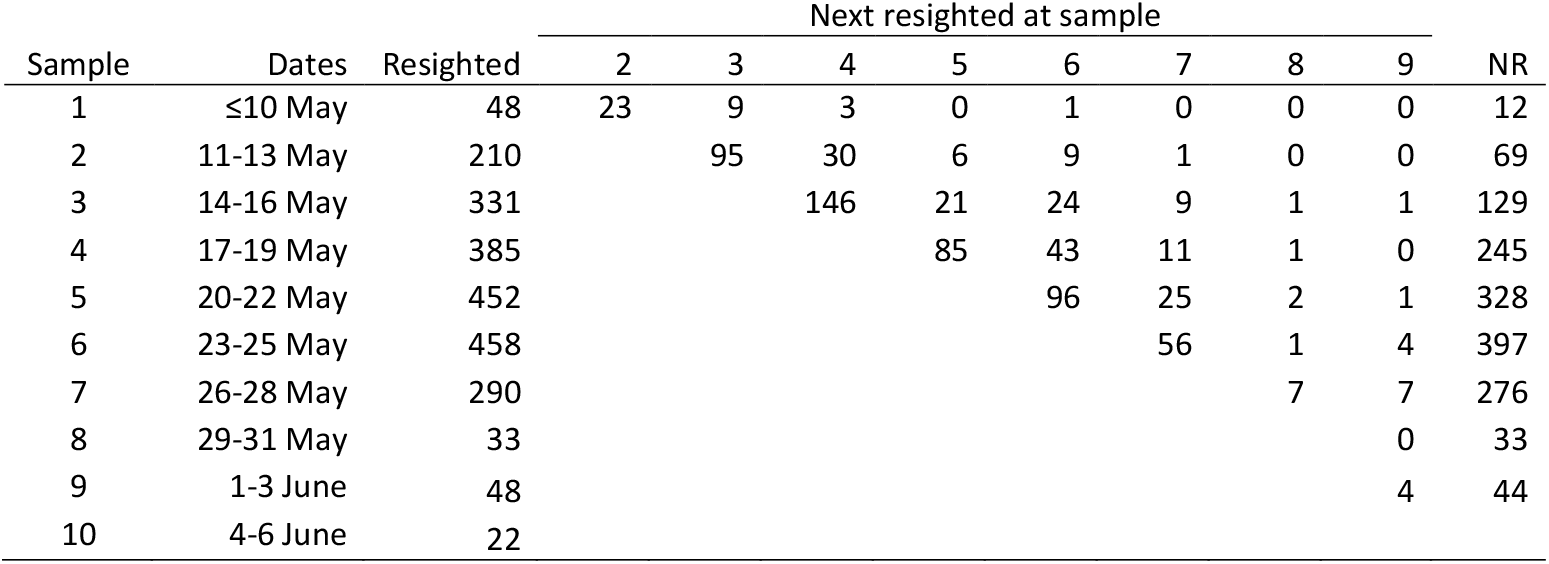

| Sample | Dates | Resighted | Next resighted at sample |  |  |  |  |  |  |  | NR |
| --- | --- | --- | --- | --- | --- | --- | --- | --- | --- | --- | --- |
|  |  |  | 2 | 3 | 4 | 5 | 6 | 7 | 8 | 9 |  |
| 1 | ≤10 May | 48 | 23 | 9 | 3 | 0 | 1 | 0 | 0 | 0 | 12 |
| 2 | 11-13 May | 210 |  | 95 | 30 | 6 | 9 | 1 | 0 | 0 | 69 |
| 3 | 14-16 May | 331 |  |  | 146 | 21 | 24 | 9 | 1 | 1 | 129 |
| 4 | 17-19 May | 385 |  |  |  | 85 | 43 | 11 | 1 | 0 | 245 |
| 5 | 20-22 May | 452 |  |  |  |  | 96 | 25 | 2 | 1 | 328 |
| 6 | 23-25 May | 458 |  |  |  |  |  | 56 | 1 | 4 | 397 |
| 7 | 26-28 May | 290 |  |  |  |  |  |  | 7 | 7 | 276 |
| 8 | 29-31 May | 33 |  |  |  |  |  |  |  | 0 | 33 |
| 9 | 1-3 June | 48 |  |  |  |  |  |  |  | 4 | 44 |
| 10 | 4-6 June | 22 |  |  |  |  |  |  |  |  |  |

## Appendix 2. Statistical Methods to Estimate Stopover Population Size Using Mark-Resight Data and Counts of Marked Birds

We converted the observations of marked birds into encounter histories, one for each bird, and analyzed the encounter histories with a Jolly-Seber (JS) model (Jolly 1965, Seber 1965, Crosbie and Manly 1985, Schwarz and Arnason 1996). The JS model includes parameters for recruitment (β), survival (ϕ), and capture (*p*) probabilities; in the context of a mark-resight study at a migration stopover site, these parameters are interpreted as probability of arrival to the study area, stopover persistence, and resighting, respectively. Stopover persistence is defined as the probability that a bird present at time *t* remains at the study area until time *t* + 1. The Crosbie and Manley (1985) and Schwarz and Arnason (1996) formulation of the JS model also includes a parameter for superpopulation size, which in our approach to mark-resight inferences for stopover populations is an estimate of the marked (leg-flagged) population size.

We chose to use 3-day periods rather than days as the sampling interval for the JS model given logistical constraints on complete sampling of the study area; multiple observations of the same individual in a given 3-day period were combined for analysis. A summary (m-array) of the mark-resight data is presented in Appendix 1.

We made inference from a fully-time dependent model; arrival, persistence, and resight probabilities were allowed to vary with sampling period [β*_t_* ϕ_*t*_ p_*t*_]. In this model, we set *p_1_* = *p_2_* and *p_k-1_* = *p_K_* (where *K* is the number of samples) because not all parameters are estimable in the fully-time dependent model (Jolly 1965, Seber 1965, Crosbie and Manly 1985, Schwarz and Arnason 1996).

We followed the methods of Royle and Dorazio (2008) and Kéry and Schaub (2012, Chapter 10) to fit the JS model using the restricted occupancy formulation. Royle and Dorazio (2008) use a state-space formulation of the JS model with parameter-expanded data augmentation. For parameter-expanded data augmentation, we augmented the observed encounter histories with all-zero encounter histories (n = 2000) representing potential recruits that were not detected (Royle and Dorazio 2012). We followed Lyons et al. (2016) to combine the JS model with a binomial model for the counts of marked and unmarked birds in an integrated Bayesian analysis. Briefly, the counts of marked birds (*m_s_*) in the scan samples are modeled as a binomial random variable:

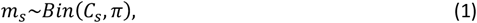

where *m_s_* is the number of marked birds in scan sample *s*, *C_s_* is the number of birds checked for marks in scan sample *s*, and π is the proportion of the population that is marked. Total stopover population size 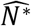 is estimated by

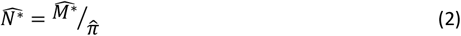

where 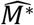 is the estimate of marked birds from the J-S model and 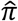 is the proportion of the population that is marked (from Eq. 1). Estimates of marked subpopulation sizes at each resighting occasion *t* 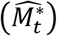 are available as derived parameters in the analysis. We calculated an estimate of population size at each mark-resight sampling occasion 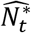 using 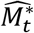 and 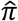 as in equation 2.

To better account for the random nature of the arrival of marked birds and addition of new marks during the season, we used a time-specific model for proportion with marks in place of equation 1 above:

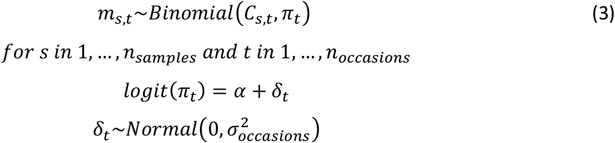

where *m_s_* is the number of marked birds in scan sample *s*, *C_s_* is the number of birds checked for marks in scan sample *s*, *δ_t_* is a random effect time of sample *s*, and *π_t_* is the time-specific proportion of the population that is marked. Total stopover population size 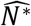 was estimated by summing time-specific arrivals of marked birds to the stopover (*B_t_*) and expanding to include unmarked birds using estimates of proportion marked:

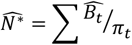

Time-specific arrivals of marked birds are estimated from the Jolly-Seber model using 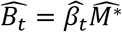 where 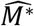 is the estimate of the number of marked birds and 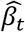 is the fraction of the population arriving at time *t*.

## Appendix 3. Number of marked-ratio scan samples

**Figure A3.1.**
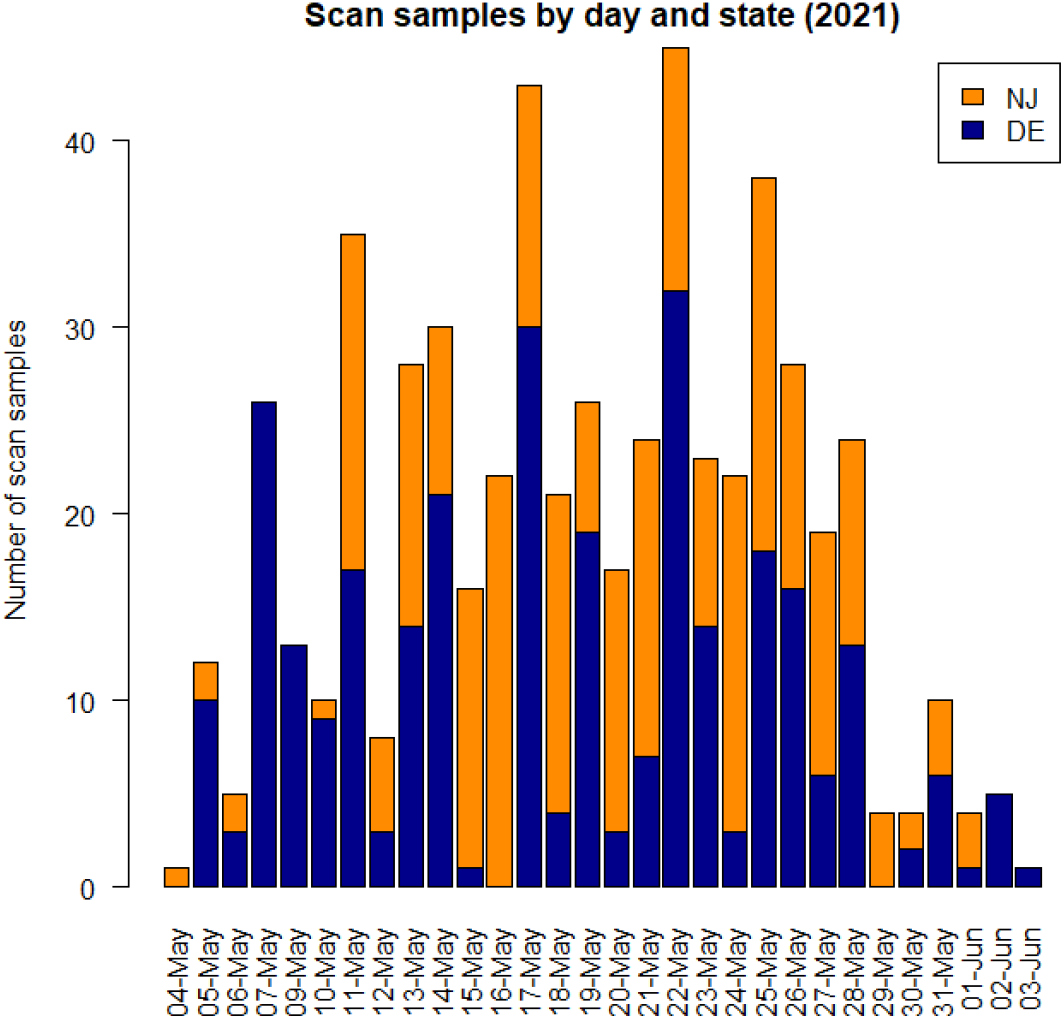
Number of marked-ratio scan samples (n = 564) collected in Delaware Bay in 2021 by field crews in Delaware (blue) and New Jersey (orange) and date. In 2021, observers in Delaware and New Jersey collected 297 and 267 scan samples, respectively.

